# Noise-induced temporary threshold shift in macaques disrupts electrophysiological temporal processing despite recovery of cochlear sensitivity and preserved ribbon synapse counts

**DOI:** 10.64898/2026.08.17.744898

**Authors:** A.N. Conner, J.A. Mondul, S. Kulkarni, C.A. Mackey, A. Batchu, N. Temghare, T.A. Hackett, R. Ramachandran

## Abstract

Noise exposure can produce lasting auditory dysfunction in the absence of permanent threshold shifts or hair cell loss, yet the functional consequences of temporary threshold shift (TTS) remain poorly defined in translational models. We assessed auditory brainstem responses (ABRs) and distortion product otoacoustic emissions (DPOAEs) in rhesus macaques (*n* = 13) at 2 and 9–10 months following a single moderate noise exposure that induced TTS. Previous histological analyses of these macaques showed no significant loss of hair cells or ribbon synapses but revealed persistent broadening of inner and outer hair cell ribbon-volume distributions. After exposure, DPOAE amplitudes and thresholds and ABR thresholds returned to pre-exposure values and showed low-frequency enhancement at later time points. Suprathreshold click- and tone-evoked ABR amplitudes were largely preserved or enhanced after exposure, consistent with compensatory gain. In contrast, macaque-specific chirp-evoked ABRs showed modest amplitude reductions and latency prolongation across waves, indicating altered neural synchrony at standard stimulus presentation rates, but with variable time courses. More temporally demanding paradigms revealed persistent impairments. ABRs to faster click rates and shorter paired-click intervals showed reduced adaptability in response amplitude and timing after normalization, with deficits persisting through 9–10 months. Increased inner hair cell ribbon-volume variability was more consistently associated with temporal response measures, including latency, paired-click recovery, and rate adaptation, than with amplitude-based ABR measures. Together, these findings reveal a lasting dissociation between response magnitude and fidelity after TTS: suprathreshold responses may be preserved or enhanced, while neural synchrony and temporal adaptability remain impaired. Increased presynaptic ribbon volume variability may serve as a structural marker of synaptic remodeling accompanying hidden auditory dysfunction, rather than as a direct determinant of suprathreshold response magnitude. Temporally demanding ABR paradigms may supplement threshold-based diagnostics for detecting persistent noise-induced auditory dysfunction.

**HIGHLIGHTS:**

- Noise-induced TTS produced suprathreshold auditory changes through 9–10 months.
- Click- and tone-evoked ABRs remained preserved or enhanced at standard presentation rates.
- Chirp, rapid-rate click, and paired-click ABRs revealed temporal processing deficits.
- Ribbon synapse counts were preserved but showed increased ribbon-volume variability.
- Ribbon-volume variability was more strongly linked to temporal than amplitude-based measures.

## 1. INTRODUCTION

Noise exposure is a major cause of auditory dysfunction and can produce both temporary and permanent hearing problems. While classic descriptions of noise-induced hearing loss (NIHL) are characterized by elevated thresholds and outer hair cell (OHC) loss, substantial hearing difficulties can occur despite normal audiograms (Le Prell, 2020; Liberman et al., 2016). Such hidden hearing loss (HHL; Schaette & McAlpine, 2011) may be associated with cochlear synaptopathy (SYN), an inner ear pathology defined by the dysfunction of synapses between inner hair cells (IHCs) and auditory nerve fibers (ANFs) (Kujawa & Liberman, 2009; Lin et al., 2011). SYN reduces the fidelity of afferent signaling to the brain (Furman et al., 2013; Suthakar & Liberman, 2021, 2022) and may contribute to speech-in-noise difficulties, impaired temporal processing, and other suprathreshold hearing deficits despite preserved audibility (Bharadwaj et al., 2019; Bharadwaj et al., 2015; Mehraei et al., 2016). SYN histopathology is robustly demonstrated in animal models following noise-induced temporary threshold shifts (TTS) after single noise exposures (Furman et al., 2013; Kujawa & Liberman, 2009; Lin et al., 2011; Mondul et al., 2026) and aging (Kujawa & Liberman, 2015; Sergeyenko et al., 2013), and is a component of sensorineural hearing loss (SNHL) when accompanied by hair cell pathology (Fernandez et al., 2020; Valero et al., 2017). However, the prevalence and functional impact of noise-induced SYN in humans remain uncertain (Viana et al., 2015; Wu et al., 2019), in part due to the lack of reliable physiological biomarkers.

Auditory brainstem responses (ABRs) are widely used to assess cochlear and brainstem function in preclinical and clinical contexts. In multiple rodent species, including mice, rats, guinea pigs, and chinchillas, SYN consistently reduces suprathreshold ABR Wave I amplitudes despite normal thresholds (Hickman et al., 2018; Kujawa & Liberman, 2009, 2015; Lin et al., 2011; Lobarinas et al., 2017; Rüttiger et al., 2013). These reductions are often accompanied by enhanced or preserved Wave V amplitudes, resulting in a lower Wave I/V ratio, which is interpreted as central gain compensation (Chambers et al., 2016; Parthasarathy & Kujawa, 2018). In humans, however, evidence for reduced Wave I amplitudes and altered I/V ratios is inconsistent: some studies report clear reductions in noise-exposed listeners with normal audiograms (Bramhall et al., 2017; Liberman et al., 2016; Skoe & Tufts, 2018), while others find no significant differences or cite methodological limitations (Grinn et al., 2017; Guest et al., 2018; Prendergast et al., 2017; Spankovich et al., 2017). Together, these mixed findings suggest that conventional ABR metrics may be insufficient to capture the full range of suprathreshold dysfunction following moderate noise exposure, particularly in humans.

Nonhuman primates (NHPs) provide a valuable translational model for auditory research, and rhesus macaques in particular exhibit cochlear anatomy and perceptual abilities that closely parallel those of humans (Burton et al., 2019; Hosoya, 2025; C. Mackey et al., 2021; Moody, 1994). Prior work has demonstrated clear histopathological evidence of SYN in macaque monkeys up to 2 months after noise-induced TTS (Valero et al., 2017). We recently extended this work to evaluate cochlear pathology through 10 months post-exposure in a larger mixed-sex cohort of macaques (Mondul et al., 2026). Despite minimal loss of hair cells or synapses, these animals exhibited persistent enlargement of IHC and OHC presynaptic ribbons. These anatomical changes were accompanied by normal or enhanced ABR Wave I amplitudes to suprathreshold clicks (Mondul et al., 2026) and permanent deficits in electrophysiological measures of suprathreshold binaural integration and behavioral spatial hearing tasks (Mackey et al., 2026).

In this study, we further characterized auditory function in the macaque cohort described by Mondul et al. (2026) and Mackey et al. (2026), using non-invasive physiological measures, including otoacoustic emissions (OAEs) and ABRs, at 2 and 9–10 months after a single noise exposure that induced TTS. Because macaque ABRs are small and highly variable across subjects, as in humans, linking physiological outcomes to postmortem histology is a critical step toward developing sensitive translational biomarkers of noise-induced auditory dysfunction (Ng et al., 2015; Stahl et al., 2022). We used a longitudinal design and a broad battery of standard and alternative ABR paradigms to improve sensitivity to subtle dysfunction over clinically relevant post-exposure intervals. To probe cochlear integrity, neural synchrony, and temporal adaptability following noise exposure, we incorporated temporally demanding ABR paradigms, including macaque-specific chirps to optimize synchrony (Elberling et al., 2010), rapid click trains to probe recovery dynamics (Burkard & Sims, 2001, 2002), and paired-click measures to assess adaptation and temporal resolution (Fujihira et al., 2024; Lee et al., 2021). We then related these functional measures to postmortem histological metrics of IHC and OHC survival, IHC-ANF synapse counts, OHC ribbon counts, IHC and OHC ribbon volumes, and cholinergic olivocochlear innervation density (Mondul et al., 2026). Taken together, this integrative approach aimed to clarify the physiological consequences of moderate noise-induced TTS and to refine translational biomarkers of noise-induced auditory dysfunction in a primate model.

## 2. MATERIALS AND METHODS

### 2.1 Subjects

Thirteen young adult rhesus macaques (*Macaca mulatta*; *n* = 4 females, 9 males) were included (mean age at study start: 6.6 years, SD = 0.65; range: 6-8 years). Although a sex-balanced cohort was sought, NHP availability constraints due to the COVID-19 pandemic limited enrollment. This age range corresponds to ∼18-30 human years (Davis & Leathers, 1985), and age-related hearing loss was not expected (Ng et al., 2015). All animals underwent baseline auditory testing, a noise exposure to induce temporary threshold shift (TTS), and post-exposure testing at 2 months (*n* = 13). Ten of these subjects also underwent long-term follow-up at 8-10 months (DPOAEs: *n* = 10; ABRs: *n* = 6). Animals were randomly assigned to study endpoints at either 2 months (*n* = 3) or 8-10 months (*n* = 10) post-exposure.

Monkeys were obtained from the California National Primate Research Center (UC Davis) and the Oregon National Primate Research Center (OHSU) and housed at the Vanderbilt University Medical Center (VUMC) on a 12:12-hour light:dark cycle. All animals received a commercial primate diet (LabDiet 5037 or 5050) supplemented with produce and foraging items, and rotated environmental enrichment (manipulanda, auditory, visual, olfactory). Filtered water was provided daily, but access was regulated due to participation in parallel behavioral studies.

Cranial implants were maintained for head fixation in those studies. All procedures were approved by the Institutional Animal Care and Use Committee (IACUC) at VUMC and strictly complied with the National Institutes of Health (NIH) guidelines.

### 2.2 Noise exposure and auditory physiological characterization

#### 2.2.1 Anesthesia

Animals were anesthetized for auditory physiological testing and the noise exposure procedure. Initial sedation was induced with an intramuscular injection of ketamine (10 mg/kg) and midazolam (0.05 mg/kg), followed by treatment with atropine or glycopyrrolate to minimize mucous secretions. Animals were intubated, and anesthesia was maintained with isoflurane (1-2%). Although anesthetic state, including isoflurane anesthesia, can influence ABR thresholds, amplitudes, and latencies (Bielefeld, 2014; Ruebhausen et al., 2012; Stronks et al., 2010), all ABR comparisons were made within subjects across pre- and post-exposure time points using the same anesthetic protocol. All anesthetized procedures were conducted in a sound-treated booth (Acoustic Systems, Austin, TX). Subjects were monitored intensively for a minimum of 72 hours post-procedure. Physiological testing was conducted 1-3 months prior to noise exposure, immediately after noise exposure (DPOAEs only, same session), 2 months post-exposure, and 9-10 months post-exposure.

#### 2.2.1 Noise exposures

Subjects underwent a single noise exposure intended to cause TTS. The noise exposure procedure was described in Mondul et al. (2026) and was similar to those previously reported by our laboratory (Burton et al., 2020; Hauser et al., 2018; C. A. Mackey et al., 2021; Valero et al., 2017). The subject was lying prone on a table with the head slightly elevated in a sound-treated booth. Closed-field speakers (MF1, Tucker-Davis Technologies) were coupled to the ears using 1.5” PE tubing and pediatric ER-3A insert earphones that were deeply inserted into each ear canal. Octave-band noise (2–4 kHz) was presented simultaneously to both ears at 120 dB SPL for four hours. The level of the exposure stimulus varied by less than 0.3 dB SPL throughout the four-hour procedure.

#### 2.2.2 Distortion product otoacoustic emission (DPOAE) testing

A Scout Biologic OAE System (Natus) was used to measure DPOAEs in response to two primary tones, *f*_1_ and *f*_2_, at eight frequencies per octave from *f*_2_= 0.5 kHz to *f*_2_= 10 kHz with a frequency ratio of *f*_2_/ *f*_1_ = 1.22 and a level ratio of L_2_ = L_1_-10 dB SPL from L_2_= 20 to L_2_= 70 dB SPL in 5-dB steps. DPOAEs (2*f*_1_-*f*_2_) collected at L_1_/ L_2_= 65/55 were used to plot DP-grams, and DPOAEs collected at increasing L_2_ levels were used to derive threshold-versus-frequency and input-output functions. *Threshold* was defined as the lowest *f*_2_ level required to produce a DPOAE with amplitude at least 0 dB SPL and at least 6 dB SPL above the noise floor. In a small subset of subjects (n= 3 at 2 months post-exposure, n= 6 at 8-10 months post-exposure), a high-frequency DPOAE system (ER-10X, Interacoustics) was used to examine responses for *f*_2_= 1-32 kHz; all other stimulus parameters and DPOAE measures were the same.

#### 2.2.3 Auditory brainstem response (ABR) testing

ABR recording methods overlapped with previous publications from our laboratory (Hauser et al., 2018; Stahl et al., 2022; Valero et al., 2017). Subdermal needle electrodes (Rhythmlink, Columbia, SC) were placed on the mastoid (active), vertex (reference), and shoulder (ground), and connected to a Medusa 4Z preamplifier (Tucker-Davis Technologies, Alachua, FL). Electrode impedances were maintained below 1 kΩ. Acoustic stimuli were delivered via closed-field speakers (MF1, Tucker-Davis Technologies) coupled to the ear with pediatric ER-3A foam tips.

Stimuli were generated in SigGenRZ and presented using an RZ6 Multi-I/O Processor (Tucker-Davis Technologies). Stimuli included broadband clicks (0.001-97 kHz, 100 µs), frequency-specific tone bursts (0.5-32 kHz), macaque-specific chirps (0.5-32 kHz, 1.6 ms), and paired clicks with varying inter-click intervals (ICIs). Clicks and tone bursts were presented at 27.7/s across 90-10 dB SPL in 5-to 10-dB steps using alternating polarity. Additional click trains were presented at increasing rates (57.7, 100, 125, 166.6, and 200/s at 70-90 dB SPL), and click pairs were presented at ICIs of 1, 2, 4, 8, and 10 ms. Stimuli were calibrated (±1 dB SPL) using a 0.5 cc coupler and verified in the ear canal with a probe-microphone system (Fonix 8000, Frye).

Signal acquisition and online filtering (300-3000 Hz) were performed in BioSigRZ. Artifact rejection excluded responses > ±1 mV. Two artifact-free waveforms (1024 repetitions each, 2048 total) were averaged, inverted, and low-pass filtered at 1.5 kHz to isolate the ABR waveform, consistent with the known spectral limits of the ABR signal (Boston, 1981)

ABR thresholds and waveform metrics were quantified as follows. For each stimulus frequency and level, the ABR threshold was defined as the lowest level (nearest 5 dB SPL) that elicited a visually identifiable waveform exceeding the noise floor (≥ 20-40 nV). Peak-to-trough amplitudes and absolute latencies were manually identified for Waves I, II, and IV. For chirp stimuli, the chirp duration (1.6 ms) was subtracted from measured latencies to account for the delay between stimulus onset and peak acoustic energy. Wave I, II, and IV input-output functions were constructed from amplitude values across stimulus levels.

For click trains at increasing presentation rates, amplitudes of Waves I, II, and IV at each rate were normalized to those at the lowest rate (27.7/s) within the same recording session to evaluate rate-dependent adaptation. These normalized amplitude values were used to construct adaptation functions across stimulus rates. Latencies for Waves I, II, and IV were baseline-subtracted by subtracting the latency measured at the lowest rate (27.7/s), yielding rate-dependent latency shifts independent of absolute latency differences across ears.

For paired-click analyses, responses were measured at ICIs of 1, 2, 4, 8, and 10 ms to assess short-term neural adaptation and recovery. For shorter ICIs in which Click 1 and Click 2 responses overlapped (1, 2, and 4 ms), the Click 1 response was subtracted from the full waveform trace to isolate the evoked response to Click 2 (Stahl et al., 2022). ICI50 was calculated from raw Wave II Click 2 amplitudes as a summary metric of short-term neural recovery. For group-level comparisons, ICI50 was derived from the selected linear mixed-effects model, which modeled raw Click 2 amplitude as a quadratic function of ICI, with time point, stimulus level, ear, and subject included. Models were fit using maximum likelihood and compared using BIC. For each time point, ICI50 was defined as the ICI at which the model-predicted Click 2 amplitude reached the midpoint between the predicted amplitudes at 1 and 10 ms ICI at 80 dB SPL. Confidence intervals and pairwise time point comparisons were estimated using a parametric bootstrap.

Because structure-function analyses required one recovery value for each ear, ear-level ICI50 values were calculated separately from raw Wave II Click 2 amplitudes using linear interpolation across the measured ICI values. These ear-level values were used to compute correlations with postmortem histological measures, whereas group-level time-point effects were tested using the mixed-effects model-derived ICI50 estimates. To evaluate response suppression independent of raw amplitude differences, Wave I, II, and IV Click 2 amplitudes were normalized to the corresponding Click 1 amplitudes at the same level. Click 2 latencies for Waves I, II, and IV were baseline-subtracted by subtracting the corresponding Click 1 latency within the same waveform, yielding ICI-dependent latency shifts independent of absolute latency.

### 2.3 Cochlear histology and quantification

Cochlear tissue collection and processing, immunohistochemistry, confocal imaging, and image quantification have been previously described (Burton et al., 2020; Mondul et al., 2026; Valero et al., 2017). Briefly, following completion of physiological testing, animals were euthanized via intravenous overdose of sodium pentobarbital and sodium phenytoin (Euthasol; 130 mg/kg), followed by transcardial perfusion with 0.9% phosphate-buffered saline (PBS) and 4% phosphate-buffered paraformaldehyde (PFA). Temporal bones were extracted, the round and oval windows were opened, and cochleae were perfused through the scala tympani with 4% PFA, then immersed in fixative for 2 hours at room temperature. Cochleae were subsequently decalcified in 0.12 M EDTA at 4°C.

Decalcified cochleae were dissected into whole mounts of the organ of Corti and immunolabeled to visualize hair cells (myosin VIIa), presynaptic ribbons (CtBP2), postsynaptic glutamate receptor patches (GluA2), and cholinergic efferent fibers (ChAT). Confocal z-stack images were obtained at half-octave-spaced locations along the cochlear spiral corresponding to frequency regions from 0.125 to 32 kHz. The following cochlear anatomical indices were quantified at each frequency place: OHC survival, IHC survival, afferent IHC-ANF synapse counts (co-localized presynaptic ribbon and postsynaptic glutamate receptor puncta), OHC ribbon counts, IHC and OHC ribbon volumes, and cholinergic efferent innervation in the OHC and IHC regions to estimate medial and lateral olivocochlear innervation density, respectively. Comprehensive cochlear histopathological characterization of this cohort is detailed in Mondul et al. (2026)

### 2.4 Statistical analyses

#### 2.4.1 Software and modeling framework

All statistical analyses were performed in MATLAB R2026a (MathWorks, Natick, MA) and Python 3.10 (Python Software Foundation) using the Statistics and Machine Learning Toolbox in MATLAB and standard scientific computing libraries in Python (NumPy, pandas, statsmodels, and SciPy). Linear mixed-effects (LME) models for the primary auditory physiology analyses were fit in MATLAB using maximum likelihood (ML). Structure–function mixed-effects models implemented in Python were fit using restricted maximum likelihood (REML). Ordinary least-squares regression with subject-clustered standard errors and Spearman rank correlations were also performed in Python.

#### 2.4.2 Model specification for auditory physiology, cochlear histology, and structure–function analyses

LME models were used for auditory outcomes with repeated measurements within subjects or subject–ear units, including DPOAE measures, ABR thresholds, and ABR amplitudes and latencies across click, chirp, rapid-rate click, and paired-click paradigms. Time point was modeled categorically, with levels corresponding to baseline, immediately post-exposure, 2 months, and 9–10 months, as applicable. Models included at least a random intercept for subject, with subject-by-ear intercepts and random slopes added when supported by model fit and diagnostics. Ear was included as a fixed effect when needed to account for systematic lateral differences.

Predictor coding depended on the outcome. Frequency was modeled continuously with linear and quadratic terms for DPOAE amplitude and threshold analyses and categorically for tone-evoked ABR threshold, amplitude, and latency analyses. Continuous frequency terms were mean-centered before calculating quadratic effects (Schielzeth, 2010). Stimulus level, presentation rate, and interclick interval were modeled continuously and were centered or z-scored when noted. Interaction and quadratic terms were included when required to test prespecified hypotheses or supported by model comparison.

Histological analyses followed Mondul et al. (2026). Hair cell survival, synapse and ribbon counts, ribbon volume, and efferent innervation were analyzed using mixed-effects models incorporating group, cochlear frequency, subject, and ear. Ribbon-volume distributions were compared using two-sample Kolmogorov–Smirnov tests with Benjamini–Hochberg correction, and ribbon-volume ranges were estimated using a jackknife procedure.

For structure–function analyses, mixed-effects models with a random intercept for subject–ear were used when outcomes contained repeated frequency-specific observations. Ordinary least-squares regression with subject-clustered standard errors was used for broadband or averaged outcomes with one observation per subject–ear, including click- and chirp-evoked ABRs, Wave II ICI50, and rate-adaptation slope. Spearman rank correlations were used as sensitivity analyses.

#### 2.4.3 Model selection and inference

For auditory-physiology analyses, candidate model structures were evaluated using Akaike and Bayesian information criterion, residual diagnostics, convergence behavior, and biological interpretability. Structure–function models were specified according to the data structure and the biological hypothesis being tested. Hypothesis-driven interaction terms were retained when needed to evaluate time-dependent or frequency-dependent relationships, regardless of statistical significance. Quadratic predictor terms were retained when they improved model fit and were supported by diagnostic evaluation.

Fixed-effect coefficients are reported with standard errors and associated two-sided hypothesis tests using a significance threshold of α = 0.05. Where reported, 95% confidence intervals were calculated from the fitted-model covariance matrix. A significant association at one time point and a nonsignificant association at another were not interpreted as evidence of a time-dependent difference unless the corresponding interaction term was significant. Simple slopes were estimated from interaction models when needed to characterize time point- or frequency-specific relationships.

#### 2.4.4 Model diagnostics

Model assumptions were evaluated by visually inspecting residual-versus-fitted plots and Q–Q plots to assess homoscedasticity and approximate normality of the residuals. Model convergence, random-effects variance estimates, and covariance matrix stability were also examined. Models that produced unstable or unreliable estimates were not used in the reported results.

#### 2.4.5. Post hoc testing and multiple comparisons

Post hoc contrasts were used to examine significant main effects and interactions in greater detail. Estimated marginal differences, simple slopes, standard errors, confidence intervals, and hypothesis tests were calculated from the fixed-effect estimates and their covariance matrix. For interactions involving time point, slopes were estimated separately at each post-exposure interval. For interactions involving frequency bin, slopes were estimated within the low-, mid-, and high-frequency regions, and differences between slopes were tested relative to the reference bin. Where multiple post hoc comparisons were performed, *p*-values were adjusted using the Benjamini–Hochberg false-discovery-rate procedure (Benjamini & Hochberg, 1995) within each prespecified family of related tests.

## 3. RESULTS

Young adult macaques underwent a single noise exposure intended to produce temporary threshold shift (TTS) and cochlear synaptopathy (SYN), followed by longitudinal assessment of non-invasive auditory physiology using OAEs and ABRs. OAE measures were obtained at baseline, immediately post-exposure, and at 2- and 9-10 months post-exposure, whereas ABR measures were obtained at baseline and at the 2- and 9-10-months post-exposure. We first present longitudinal physiological outcomes across threshold-based, otoacoustic, and suprathreshold ABR measures, then summarize cochlear histology, and finally evaluate structure–function relationships.

### 3.1 Peripheral auditory function is largely preserved after noise-induced temporary threshold shift

#### 3.1.1 DPOAE amplitudes recover following TTS and show low-frequency enhancement at later time points

DP-grams measured across f2 frequencies from 0.5–10 kHz at L1/L2 = 65/55 dB SPL (Fig. 1A–B) showed a large, frequency-dependent reduction in DPOAE amplitude immediately after noise exposure. Significant reductions extended from ∼0.7–10 kHz, with the largest decrease occurring near 5.4 kHz (peak Δ = −30.49 dB SPL; Table 1). By 2 months post-exposure, DPOAE amplitudes had largely recovered, with no FDR-significant frequency-specific reductions from baseline. At 9–10 months, DPOAE amplitudes remained modestly enhanced at low frequencies, with significant increases localized primarily from 500–2,000 Hz (Table 1). In the smaller subset of ears tested over an extended high-frequency range from f2 = 1–32 kHz (Fig. 1C–D), immediate post-exposure suppression was also observed, with significant reductions extending from ∼1.0–13.5 kHz. No significant changes from baseline were detected in this extended high-frequency dataset at either 2 or 9–10 months post-exposure.

**Figure 1.**
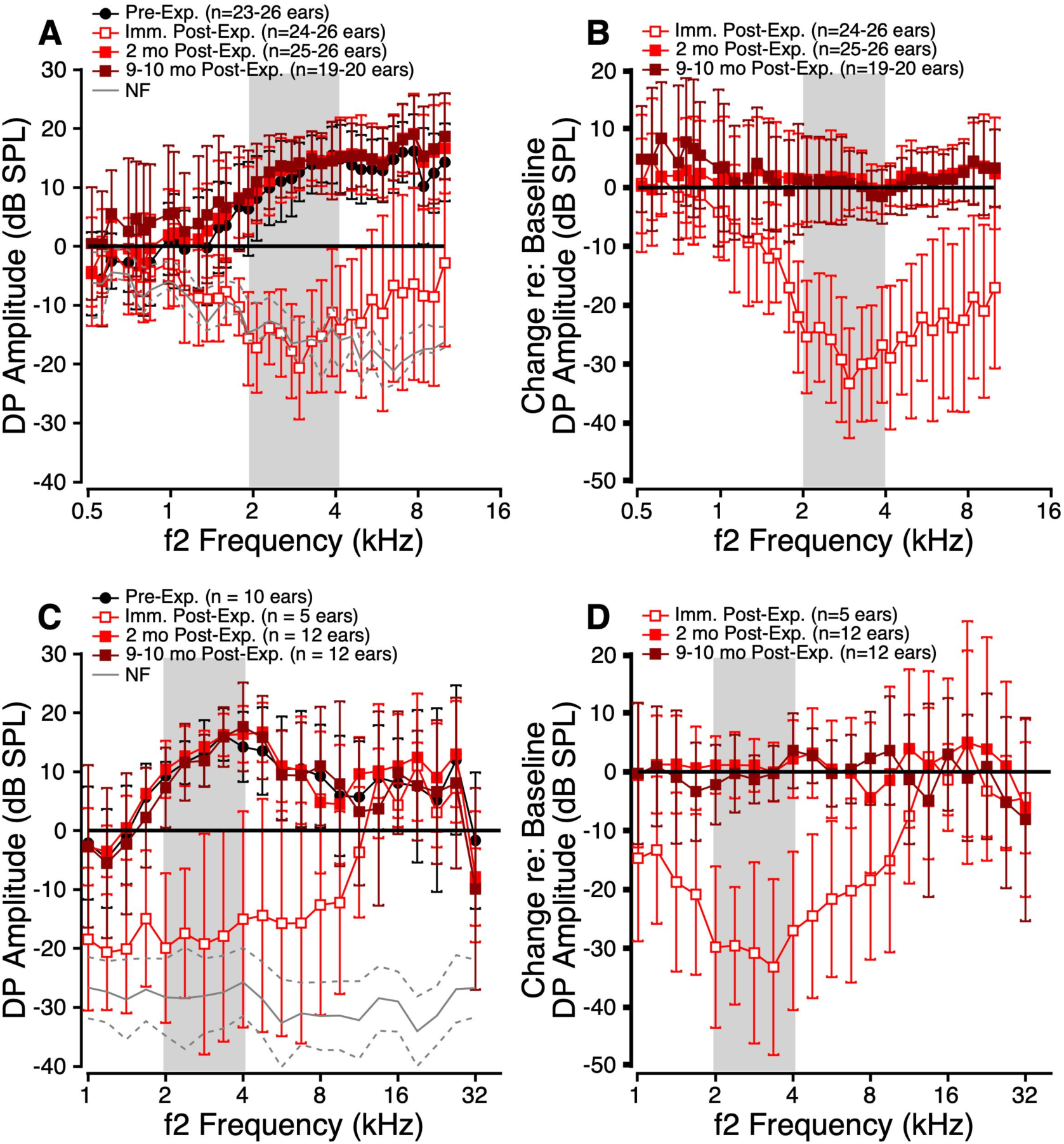
DP-grams following noise exposure. **(A, C)** Mean DPOAE amplitudes across *f*2 frequency at four time points: pre-exposure, immediately post-exposure, 2 months post-exposure, and 9–10 months post-exposure. DP-grams are shown over two frequency ranges: 0.5– 10 kHz (A) and 1–32 kHz (C). Symbols indicate pre-exposure values (solid black circles), immediately post-exposure values (open red squares), 2-month post-exposure values (solid red squares), and 9–10-month post-exposure values (solid maroon squares). Shaded regions indicate the noise exposure band. **(B, D)** Mean change in DPOAE amplitude relative to pre-exposure across time points. Error bars represent ±1 SD.

**Table 1.** Summary of post-exposure DPOAE outcomes. Consolidated summary of DPOAE findings across three complementary outcome measures: fixed-level DPOAE amplitudes (65/55 dB SPL), across-level amplitude growth functions, and DPOAE thresholds (lowest level producing a detectable DP). Results are organized by frequency band (Low: <2 kHz; Mid: 2–6 kHz; High: 7–10 kHz) and by post-exposure timepoint relative to baseline. Only statistically significant changes (FDR-corrected *p* < 0.05) are shown.

| Timepoint After Exposure | Frequency Band | Fixed-Level (65/55) DPOAE Amplitudes | Across-Level DPOAE Growth Functions | DPOAE Thresholds |
| --- | --- | --- | --- | --- |
| Immediate | Low (<2 kHz) | Reduced | Reduced (25–80 dB) | Elevated (516–844 Hz) |
|  | Mid (2–6 kHz) | Large reduction | Reduced (0–80 dB) | Elevated (516–9,141 Hz) |
|  | High (7–10 kHz) | ns | ns | Elevated (some HF bins) |
| 2 Months | Low (<2 kHz) | ns | ns | Slight reduction |
|  | Mid (2–6 kHz) | ns | ns | ns |
|  | High (7–10 kHz) | ns | ns | ns |
| 9–10 Months | Low (<2 kHz) | Enhanced | Modest enhancement | Reduced (516–1,922 Hz) |
|  | Mid (2–6 kHz) | ns | ns | ns |
|  | High (7–10 kHz) | ns | ns | ns |

DPOAE growth functions showed a similar pattern (Fig. 2). Immediately after noise exposure, amplitudes were broadly reduced, with the most robust deficits in the 2–6 kHz range. By 2 months, growth functions had largely recovered, although small residual reductions persisted at higher stimulus levels in low- and mid-frequency regions. At 9–10 months, DPOAE amplitudes were modestly higher overall, consistent with the delayed low-frequency enhancement observed in the DP-grams. Responses from 7–10 kHz did not differ significantly from baseline at any post-exposure time point.

**Figure 2.**
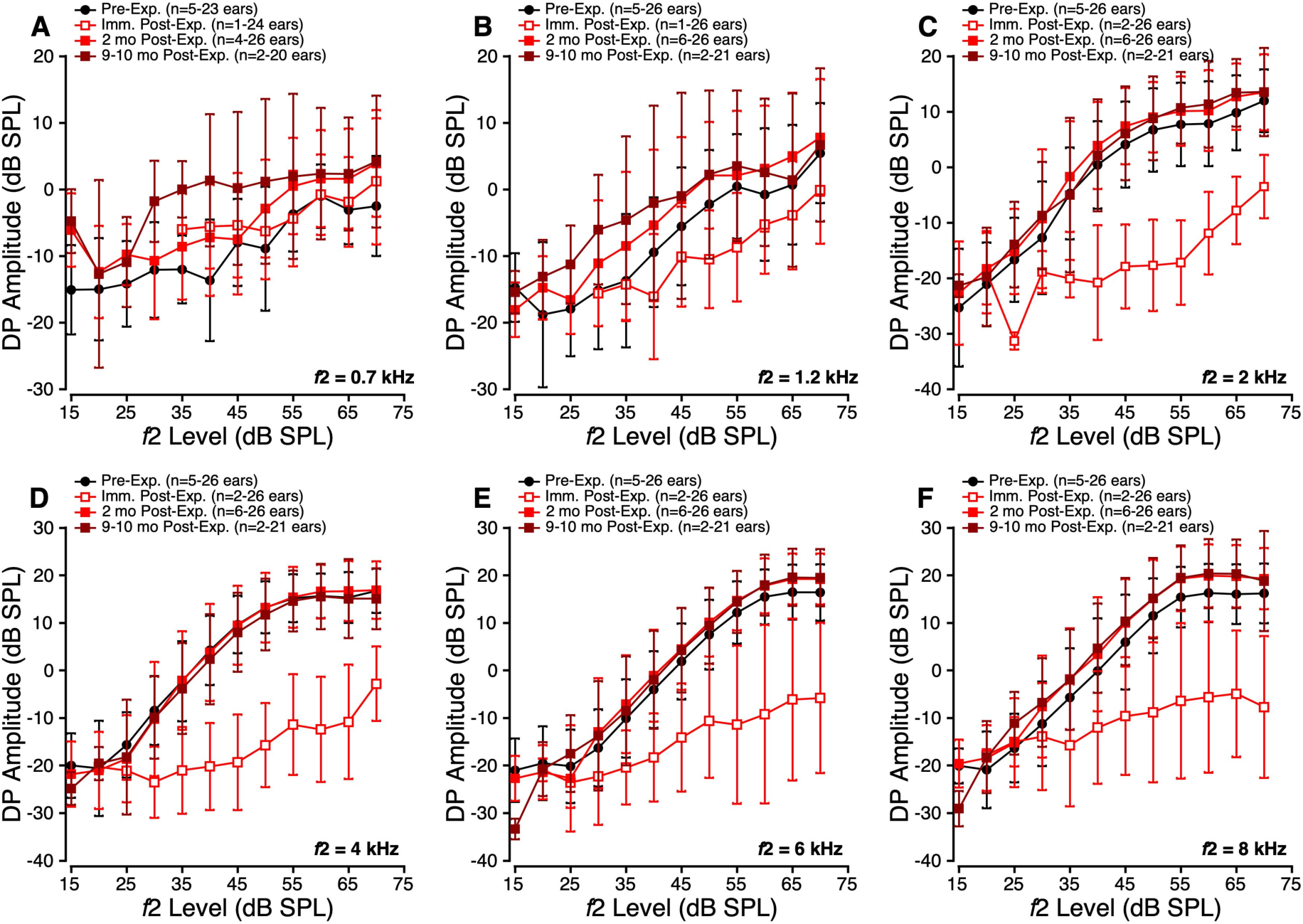
DPOAE input-output functions following noise exposure. Mean DPOAE amplitudes (dB SPL) across *f*2 levels (dB SPL) at six *f*_2_ frequencies: 0.7 kHz (A), 1.2 kHz (B), 2 kHz (C), 4 kHz (D), 6 kHz (E), 8 kHz (F) at four time points: pre-exposure (solid black circles), immediately post-exposure (unfilled red squares), 2 months post-exposure (solid red squares), and 9-10 months post-exposure (solid maroon squares). Error bars represent ±1SD. Because individual-ear DPOAE growth functions exhibit substantial variability, particularly at low frequencies and higher stimulus levels, the visually apparent differences in the raw mean curves may be smaller than the statistically significant effects detected by the mixed-effects model.

#### 3.1.2 DPOAE and ABR thresholds recover following TTS and show low-frequency enhancement at later time points

DPOAE thresholds measured across f2 frequencies from 0.5–10 kHz (Fig. 3A–B) increased immediately after noise exposure, indicating an acute threshold shift. Significant threshold elevations extended from 516–9,141 Hz, with the largest increase near 5.4 kHz (peak Δ = +32.61 dB SPL; Table 1). By 2 months post-exposure, DPOAE thresholds had recovered, with small but significant reductions relative to baseline at low frequencies from 516–844 Hz. At 9– 10 months, thresholds remained lower than baseline across a broader low-frequency range from 516–1,922 Hz, with no significant threshold elevation above ∼2 kHz.

**Figure 3.**
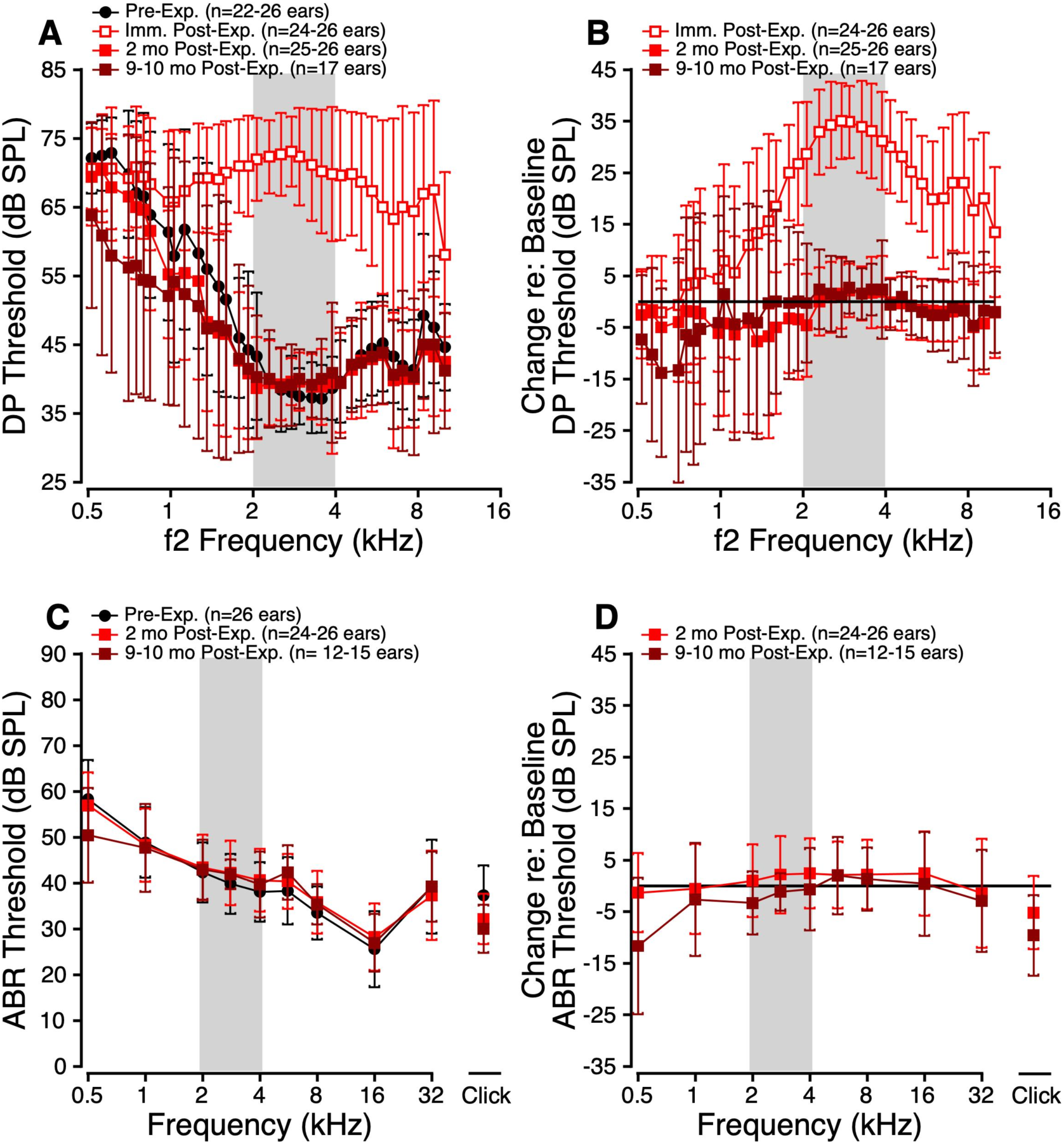
DPOAE and ABR thresholds following noise exposure. **(A)** Mean DPOAE thresholds across f2 frequency from 0.5–10 kHz at pre-exposure, immediately post-exposure, 2 months post-exposure, and 9–10 months post-exposure. Symbols indicate pre-exposure values (solid black circles), immediately post-exposure values (open red squares), 2-month post-exposure values (solid red squares), and 9–10-month post-exposure values (solid maroon squares). The shaded region indicates the noise exposure band. **(B)** Mean change in DPOAE threshold relative to pre-exposure**. (C)** Mean ABR thresholds across frequency from 0.5–32 kHz across time points**. (D)** Mean change in ABR threshold relative to pre-exposure. Error bars represent ±1 SD. Sample sizes vary across time points and frequencies because not all ears contributed data at each follow-up interval.

Because ABR testing was not performed immediately after exposure due to anesthesia-based procedural time constraints, ABR threshold analyses were limited to baseline, 2 months, and 9–10 months post-exposure. Tone-evoked ABR thresholds measured from 0.5–32 kHz showed no evidence of persistent threshold elevation following TTS (Fig. 3C–D). At 2 months, thresholds did not differ significantly from baseline at any test frequency. At 9–10 months, ABR thresholds were reduced at 0.5 kHz, paralleling the low-frequency DPOAE threshold findings, while thresholds at all other frequencies remained comparable to baseline. Click-evoked ABR thresholds were also modestly reduced relative to baseline at both post-exposure time points.

Together, DPOAE amplitudes, DPOAE thresholds, and ABR thresholds demonstrated recovery of peripheral auditory sensitivity after TTS. Thus, the suprathreshold ABR changes described below occurred in the absence of ongoing threshold loss. A summary of the key post-exposure DPOAE findings is provided in Table 1.

### 3.2 Suprathreshold ABRs at standard presentation rates are largely preserved or enhanced after TTS but vary by stimulus type

#### 3.2.1. Click-evoked ABRs show sustained suprathreshold amplitude enhancement after exposure

Following noise exposure, click-evoked ABR Wave I amplitudes (Fig. 4B) were larger than baseline at both 2 and 9–10 months post-exposure. In addition, the Wave I amplitude-level function was steeper than at baseline, with post hoc analyses indicating that these differences were primarily observed at stimulus levels ≥45 dB SPL. Wave II amplitudes (Fig. 4C) showed a similar pattern, with sustained post-exposure enhancement that was most pronounced at higher stimulus levels, spanning 40–90 dB SPL at 2 months and 50–90 dB SPL at 9–10 months. In contrast, although Wave IV amplitudes (Fig. 4D) were also higher than baseline after exposure, the Wave IV amplitude-level function was not significantly altered.

**Figure 4.**
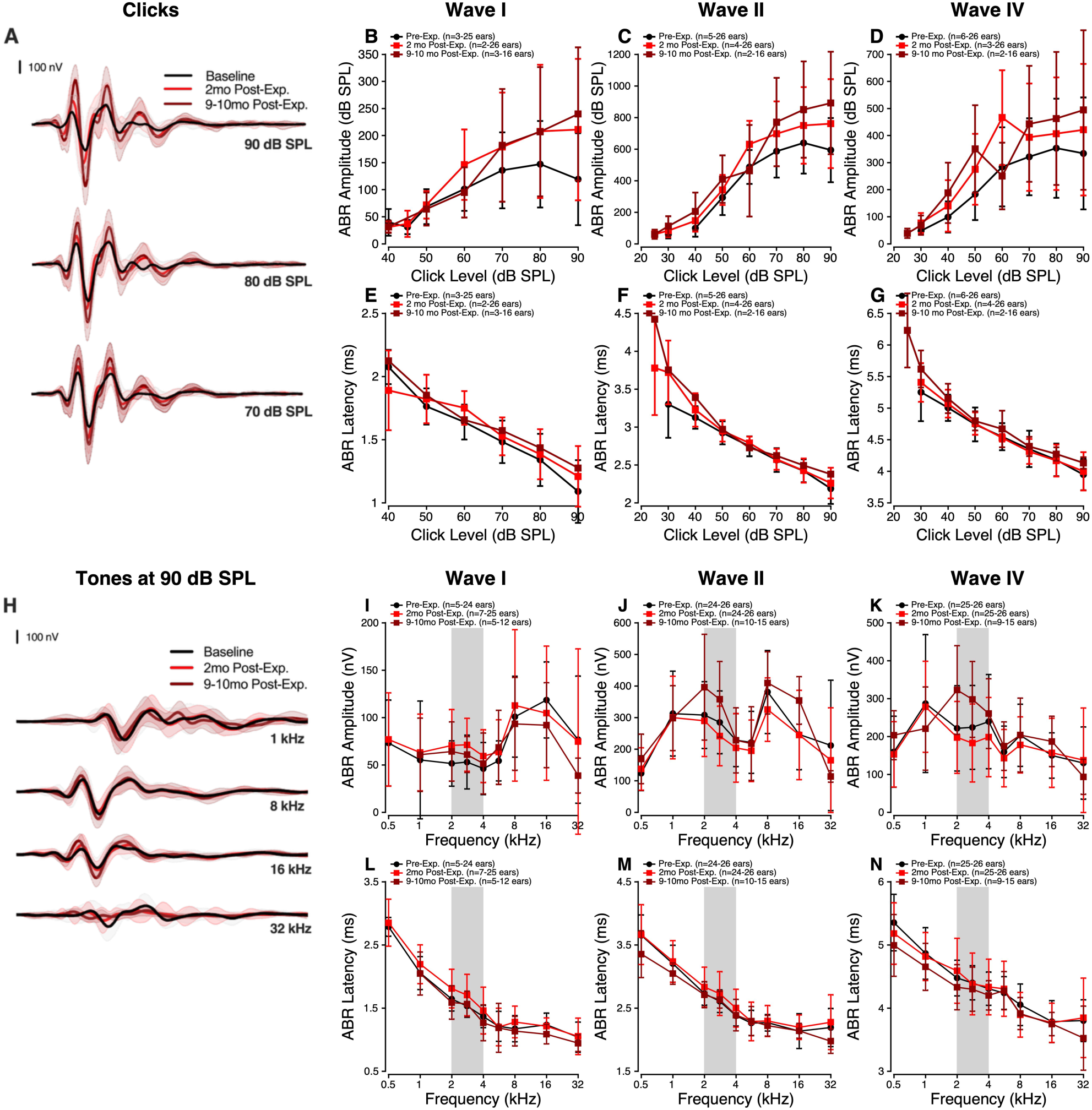
Click- and tone-evoked ABR responses after noise exposure. **(A)** Grand-average click-evoked ABR waveforms at 90, 80, and 70 dB SPL. **(B–D)** Click-evoked ABR Wave I, II, and IV amplitudes as a function of stimulus level. **(E–G)** Click-evoked ABR Wave I, II, and IV latencies as a function of stimulus level. **(H)** Grand-average tone-evoked ABR waveforms at 90 dB SPL for selected frequencies. **(I–K)** Tone-evoked ABR Wave I, II, and IV amplitudes at 90 dB SPL as a function of frequency. **(L–N)** Tone-evoked ABR Wave I, II, and IV latencies at 90 dB SPL as a function of frequency. Black circles/lines indicate baseline, red squares/lines indicate 2 months post-exposure, and dark red squares/lines indicate 9–10 months post-exposure. Shaded regions indicate the noise exposure band. Error bars indicate ±1 SD.

Click-evoked ABR Wave I latency (Fig. 4E) showed a small but significant prolongation after exposure (approximately 0.1ms). Waves II and IV (Fig. 4F–G) did not show broad mean latency shifts. Instead, latencies were prolonged only at select stimulus levels, with delays concentrated mainly at lower levels, particularly ≤40 dB SPL. Together, click-evoked ABRs remained robust after noise exposure, with enhanced suprathreshold amplitudes and modest, wave-specific latency changes.

#### 3.2.2 Tone-evoked ABRs exhibit wave- and frequency-specific changes after exposure

Because tone-evoked ABRs were measured across multiple frequencies and levels, Fig. 4H–N shows representative 90-dB SPL responses, with level-dependent effects summarized from the full input-output analyses. After noise exposure, tone-evoked ABR Wave I amplitude (Fig. 4I) was largely preserved, with no consistent difference from baseline at either post-exposure time point. At 2 months, Wave II amplitude (Fig. 4J) showed limited frequency-specific reductions rather than broad suppression, with significant decreases at 4–5.6 kHz at 80– 90 dB SPL, 8 kHz at 70–90 dB SPL, and 32 kHz at 40–80 dB SPL. By 9–10 months, Wave II amplitude was enhanced at select low- and mid-frequency regions, most clearly at 0.5 and 16 kHz, while reduced responses persisted at the highest frequency tested (32 kHz). Wave IV amplitude (Fig. 4K) was broadly reduced at 2 months but returned to baseline by 9–10 months.

Post-exposure changes in tone-evoked ABR latencies also varied by wave and frequency (Fig. 4L–N). Compared to baseline, Wave I latency (Fig. 4L) showed frequency- and level-specific changes at 2 months, with post hoc analyses across the full input-output function indicating latency prolongation at 2.8, 4, and 8 kHz and latency shortening at 1 kHz. Wave II and Wave IV latencies (Fig. 4M–N) showed stronger evidence of shortening over time, especially at low frequencies and most clearly at 9–10 months. Overall, tone-evoked ABRs did not show a simple loss of suprathreshold responsiveness, but instead varied by wave, frequency, and time after exposure.

#### 3.2.3. Chirp-evoked ABRs show modest amplitude reductions and sustained latency prolongation after exposure

Unlike click- and tone-evoked ABRs, which were largely preserved or enhanced after noise exposure, chirp-evoked ABRs showed a more consistent pattern of modest amplitude reductions and sustained latency prolongation. Wave I and Wave II amplitude reductions emerged at 9–10 months, whereas Wave IV amplitude was reduced at both post-exposure time points (Fig. 5B–D). These reductions were modest in magnitude, approximately 40–90 nV, and were not strongly level-dependent. Latencies were consistently prolonged across Waves I, II, and IV at both post-exposure time points (Fig. 5E–G), with estimated delays of approximately 0.4 ms. These latency shifts were also not strongly level-dependent, suggesting a relatively uniform delay across stimulus levels.

**Figure 5.**
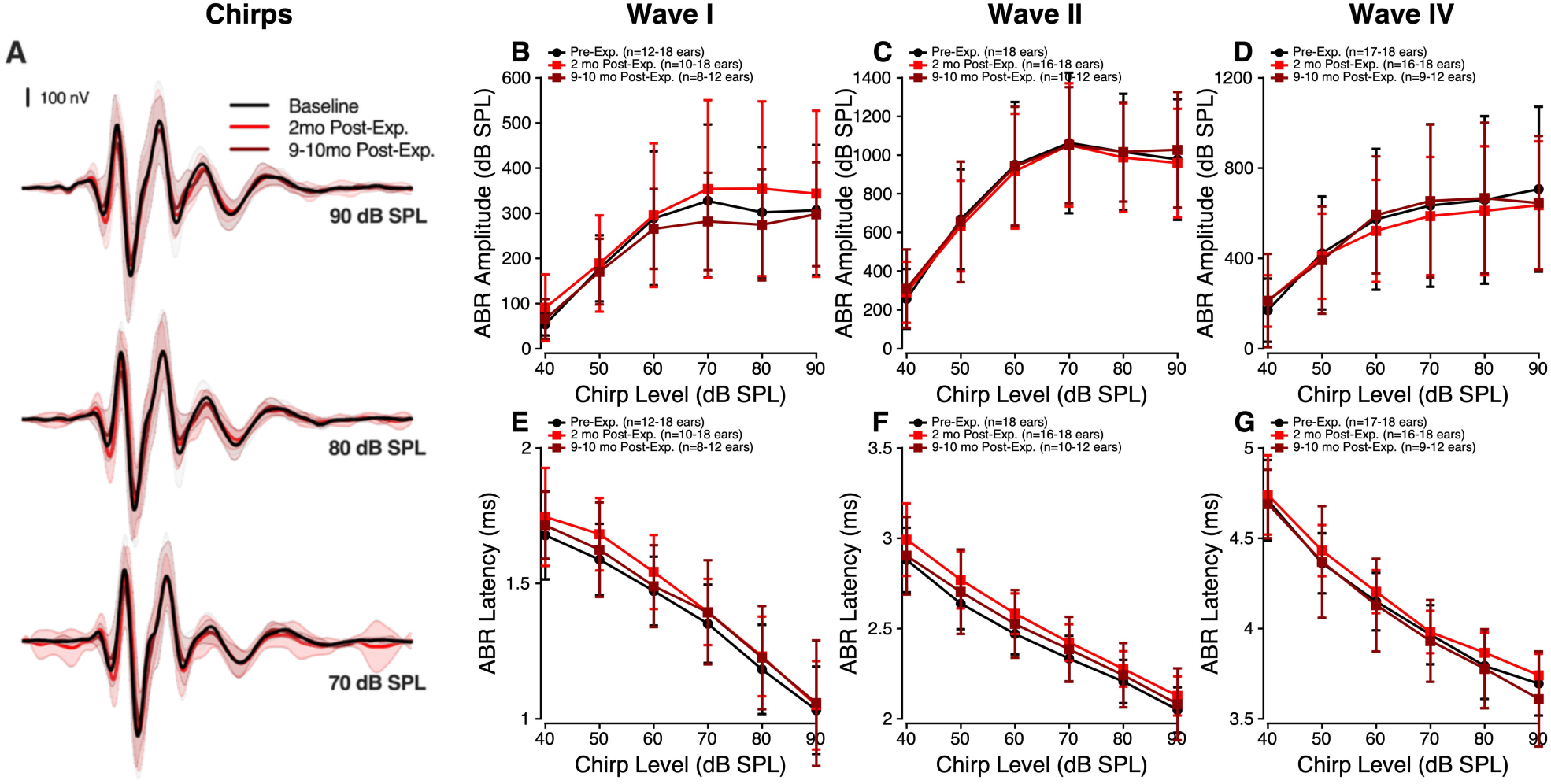
Chirp-evoked ABR responses after noise exposure. **(A)** Grand-average chirp-evoked ABR waveforms at 90, 80, and 70 dB SPL. **(B–D)** Chirp-evoked ABR Wave I, II, and IV amplitudes as a function of stimulus level. **(E–G)**, Chirp-evoked ABR Wave I, II, and IV latencies as a function of stimulus level. Black circles/lines indicate baseline, red squares/lines indicate 2 months post-exposure, and dark red squares/lines indicate 9–10 months post-exposure. Error bars indicate ±1 SD.

Together, suprathreshold ABRs obtained at standard presentation rates varied by stimulus type: click- and tone-evoked responses were largely preserved or enhanced, whereas chirp-evoked responses showed modest amplitude reductions and sustained latency delays.

### 3.3 Suprathreshold ABRs show impaired adaptation during rapid stimulation despite enhanced raw ABR amplitudes

#### 3.3.1 Rapid-rate click ABRs show enhanced raw amplitudes but reduced rate adaptation after normalization

Temporal processing was first assessed using suprathreshold clicks (70–90 dB SPL) presented across increasing stimulation rates (27.7–200 clicks/s; Fig. 6A). Raw ABR amplitudes were elevated after noise exposure across Waves I, II, and IV (Fig. 6B–D). This enhancement was broad across click rates but was most pronounced at slower presentation rates, where responses were largest. Wave II showed the strongest effect, particularly at 9–10 months post-exposure, with the largest increase observed at 27.7 clicks/s.

**Figure 6.**
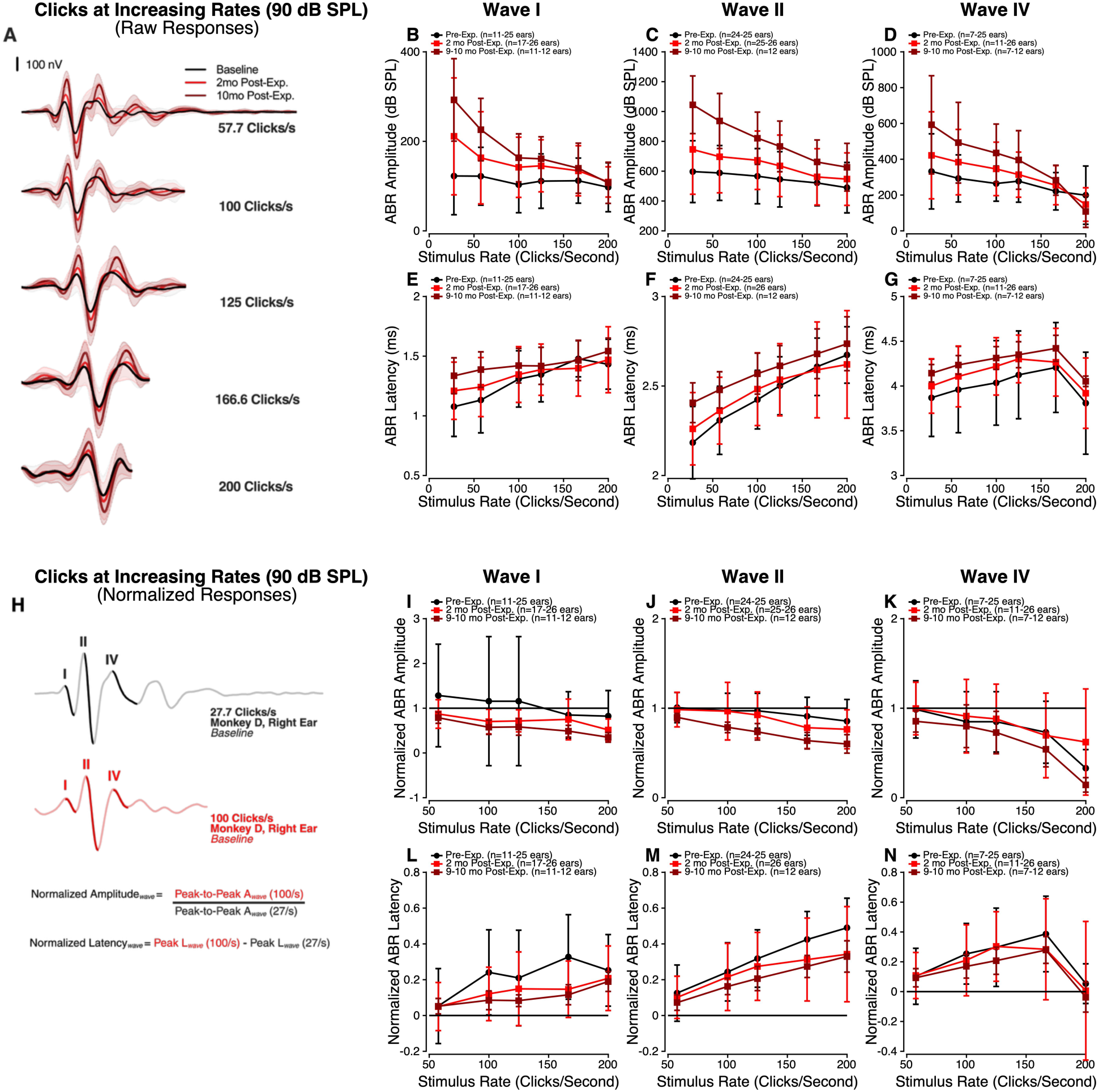
Rate-dependent suprathreshold click-evoked ABR responses after noise exposure. **(A)** Grand average ABR waveforms evoked by 90 dB SPL clicks across increasing presentation rates (27.7–200 clicks/s). **(B–D)** Mean ABR amplitudes across stimulus rate for Waves I, II, and IV. **(E–G)** Mean ABR latencies across stimulus rate for Waves I, II, and IV. **(H)** Example illustrating amplitude and latency normalization using baseline responses from one ear. Amplitudes were normalized by dividing the response at each presentation rate by the corresponding response at 27.7 clicks/s. Latencies were normalized by subtracting the latency at 27.7 clicks/s from the latency at each higher presentation rate. **(I–K)** Mean normalized ABR amplitudes across stimulus rate for Waves I, II, and IV. **(L–N)** Mean normalized ABR latencies across stimulus rate for Waves I, II, and IV. All data shown are from the 90 dB SPL condition. Statistical analyses described in the text include responses measured at 70, 80, and 90 dB SPL. Black circles: pre-exposure; red squares: 2 months post-exposure; maroon squares: 9–10 months post-exposure. Error bars = ±1 SD.

To account for differences in overall response size, amplitudes at click rates from 57.7– 200 clicks/s were normalized within each ear by dividing each response by the response at 27.7 clicks/s (Fig. 6H). After normalization, the post-exposure enhancement in raw amplitude was no longer evident. Instead, normalized amplitudes were reduced after exposure (Fig. 6I–K), with greater declines in response amplitude as click rate increased. This effect was broad across stimulation rates and was strongest at 9–10 months. Wave I and Wave II showed reductions at both post-exposure time points, whereas Wave IV showed a significant reduction only at 9–10 months.

Raw ABR latencies were prolonged after noise exposure across click rates for Waves I, II, and IV (Fig. 6E–G). To evaluate rate-dependent timing, normalized latency was calculated within each ear by subtracting the latency at 27.7 clicks/s from the latency at each higher click rate. At baseline, normalized latencies increased with click rate, as expected during more rapid stimulation. After exposure, normalized latency shifts were reduced compared to baseline for Wave I and Wave II at both post-exposure time points, whereas Wave IV did not show significant time point-dependent changes (Fig. 6L–N). Overall, raw rapid-rate click latencies were broadly prolonged after exposure, whereas latency shifts relative to 27.7 clicks/s were reduced only for Waves I and II.

#### 3.3.2 Paired-click ABRs show impaired recovery from adaptation

Suprathreshold paired-click stimuli (70–90 dB SPL) with inter-click intervals (ICIs) ranging from 10 to 1 ms were used to assess recovery from neural adaptation (Fig. 7A).

**Figure 7.**
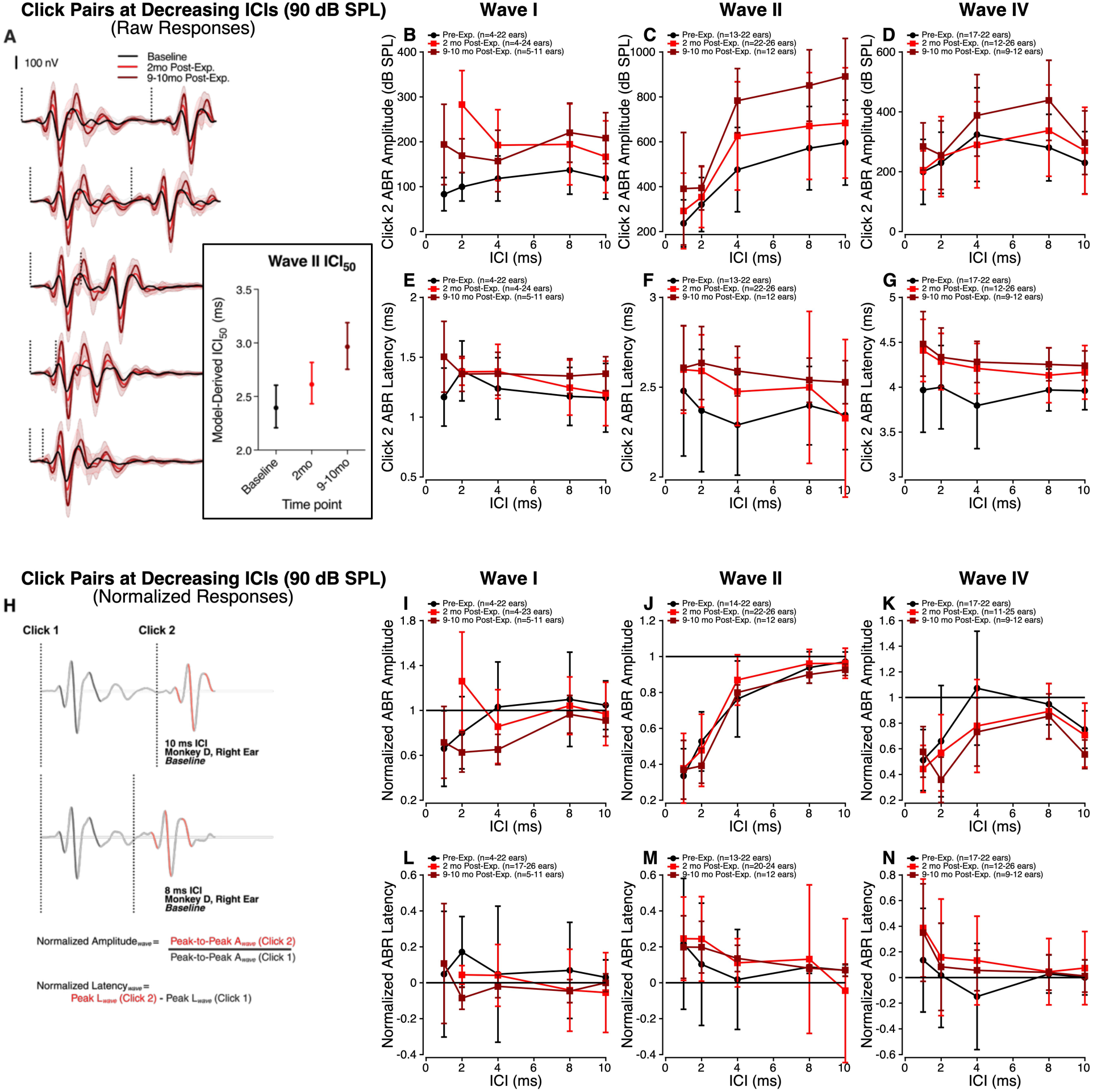
Paired-click suprathreshold ABR responses after noise exposure. **(A)** Grand average ABR waveforms evoked by 90 dB SPL click pairs across inter-click intervals (ICIs; 1– 10 ms). Inset shows model-derived Wave II ICI50 values. Inset shows model-derived Wave II ICI50 values. ICI50 was estimated from the selected mixed-effects model at 80 dB SPL and defined as the inter-click interval at which the model-predicted Click 2 amplitude reached the midpoint between the model-predicted amplitudes at 1 and 10 ms ICI. Error bars represent parametric-bootstrap 95% confidence intervals. **(B–D)** Mean Click 2 ABR amplitudes across ICI for Waves I, II, and IV. **(E–G)** Mean Click 2 ABR latencies across ICI for Waves I, II, and IV. **(H)** Example illustrating amplitude and latency normalization using baseline paired-click responses from one ear. Amplitudes were normalized by dividing the Click 2 response by the corresponding Click 1 response. Latencies were normalized by subtracting the corresponding Click 1 latency from the Click 2 latency. **(I–K)** Mean normalized ABR amplitudes across ICI for Waves I, II, and IV. **(L–N)** Mean normalized ABR latencies across ICI for Waves I, II, and IV. All data shown are from the 90 dB SPL conditions. Statistical analyses described in the text include responses measured at 70, 80, and 90 dB SPL. Black circles: pre-exposure; red squares: 2 months post-exposure; maroon squares: 9–10 months post-exposure. Error bars = ±1 SD, except where noted for ICI50.

Compared to baseline, raw amplitudes evoked by the second click (Click 2) were elevated after noise exposure across Waves I, II, and IV (Fig. 7B–D). This enhancement was broad across ICIs but was greatest at longer intervals, particularly at 9–10 months post-exposure. Wave II exhibited the largest overall enhancement.

Click 2 amplitudes were then normalized within each ear and condition by dividing the Click 2 amplitude by the corresponding Click 1 amplitude (Fig. 7H). This analysis revealed reduced paired-click recovery after exposure (Fig. 7I–K). Wave I showed a significant reduction only at 9–10 months, whereas Waves II and IV showed reductions at both post-exposure time points. Because raw Wave II amplitudes showed the largest Click 2 enhancement, recovery was further quantified by estimating Wave II ICI50, the inter-click interval required for the second-click response to reach half-maximal recovery. The mixed-effects model estimated Wave II ICI50 increased from 2.40 ms at baseline to 2.62 ms at 2 months and 2.97 ms at 9–10 months post-exposure, indicating slower recovery of Wave II responses following noise exposure (Fig. 7A, inset).

Raw Click 2 latencies showed wave- and level-specific post-exposure changes (Fig. 7E– G). Wave I latencies were prolonged at both post-exposure time points, whereas Wave II latencies showed no significant overall time point-dependent change. In contrast, Wave IV latencies were prolonged primarily at higher stimulus levels (80–90 dB SPL) at both post-exposure time points. These latency changes were not strongly dependent on inter-click interval.

Click 2 latencies were then normalized within each ear and condition by subtracting the corresponding Click 1 latency (Fig. 7L–N). Compared with normalized amplitudes, normalized latency changes were smaller and less consistent across waves. Significant shifts were observed in individual waves at selected post-exposure time points, but no common pattern of impaired latency recovery emerged. A summary of the key post-exposure ABR findings is provided in Table 2.

**Table 2.** Summary of post-exposure ABR outcomes. Consolidated summary of ABR findings across conventional and temporally demanding electrophysiological paradigms following noise-induced TTS. Results are organized by ABR assay and summarize the key post-exposure findings relative to baseline. The final column summarizes the primary physiological interpretation of each assay. Only statistically significant or biologically meaningful findings are summarized.

| **ABR Assay** | **Key Post-Exposure Finding** | **Physiological Interpretation** |
| --- | --- | --- |
| **Thresholds** | Improved click thresholds and recovered tone thresholds following TTS | Recovery and enhancement of cochlear sensitivity |
| **Click-evoked ABRs** | Enhanced suprathreshold amplitudes with modest latency changes | Enhanced suprathreshold neural responses with modest timing alterations |
| **Tone-evoked ABRs** | Mixed suprathreshold amplitude and latency changes | Preserved, heterogeneous suprathreshold neural responses |
| **Chirp-evoked ABRs** | Modest amplitude reductions and sustained latency prolongation | Reduced neural synchrony |
| **Rapid-rate click ABRs** | Enhanced raw amplitudes but reduced normalized response adaptation | Impaired temporal fidelity and neural adaptability |
| **Paired-click ABRs** | Enhanced raw Click 2 amplitudes but reduced normalized response recovery | Impaired neural recovery and temporal processing |

### 3.4 Presynaptic ribbon size variability is more consistently associated with temporal than amplitude-based suprathreshold ABR measures

#### 3.4.1 Cochlear histology reveals increased presynaptic ribbon volume variability after TTS

To identify peripheral structural changes that might underlie the suprathreshold ABR findings, cochlear histology was evaluated across multiple cellular and synaptic measures, including OHC and IHC survival, IHC synapse counts, OHC ribbon counts, cholinergic efferent innervation density, and IHC and OHC ribbon morphology. OHC and IHC survival and cholinergic efferent innervation density showed no significant group-level differences from controls and are reported in detail in Mondul et al. (2026). Because the structure–function analyses below focus on the synaptic measure that remained altered after noise exposure, Fig. 8A provides a schematic overview of the synaptic elements examined and illustrates the ribbon size variability quantified in Fig. 8D–E.

**Figure 8.**
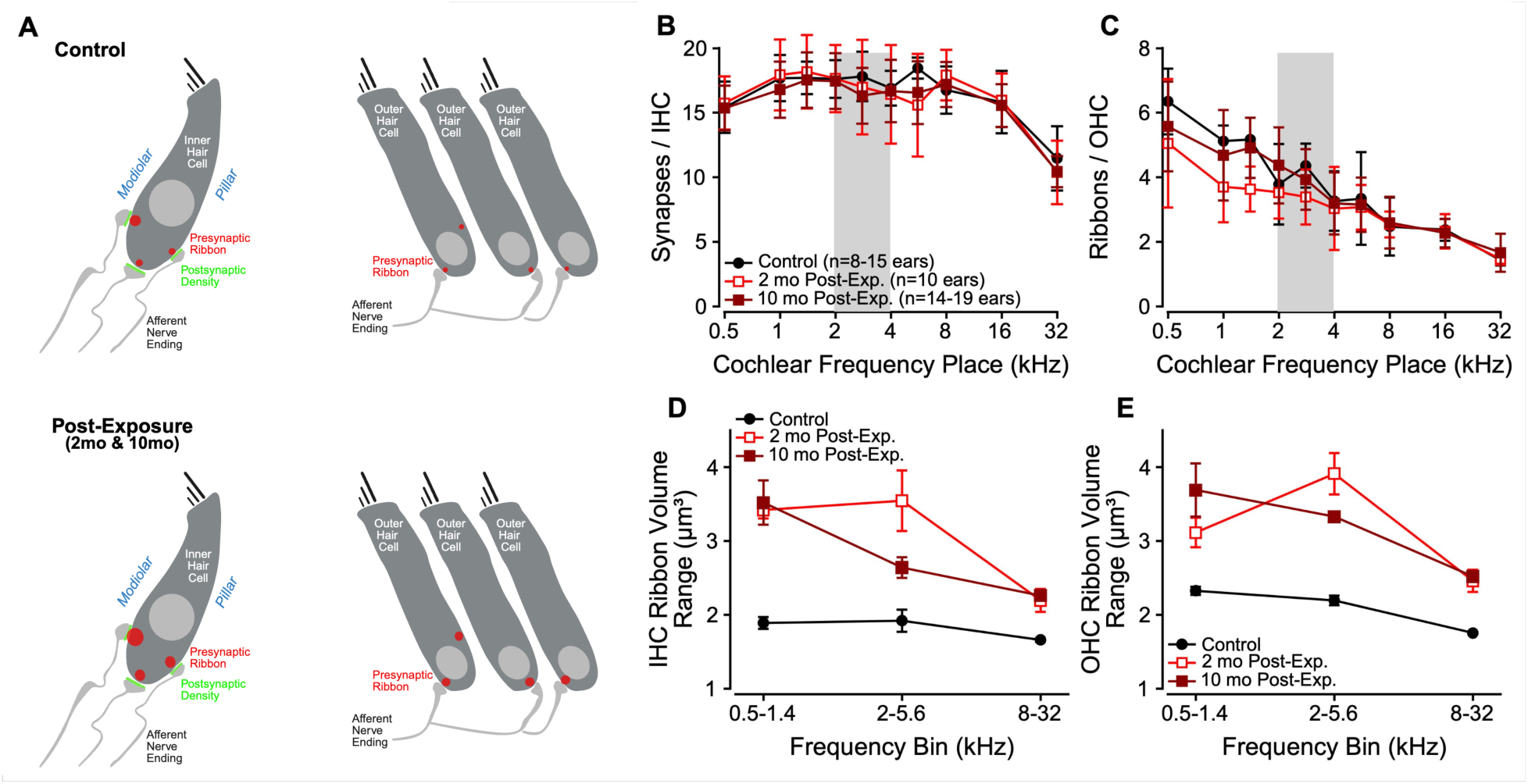
IHC and OHC synapse counts, ribbon counts, and ribbon volumes after noise exposure. **(A)** Schematic illustrating synaptic elements in control (top) and noise-exposed (bottom) cochleae, highlighting increased presynaptic ribbon volume variability despite unchanged synapse and ribbon counts following noise exposure. Presynaptic ribbons are shown in red and postsynaptic densities in green. The left panels depict inner hair cell (IHC) synapses, and the right panels depict outer hair cell (OHC) presynaptic ribbons. **(B, C)** Mean synapses per IHC **(B)** and ribbons per OHC **(C)** as a function of cochlear frequency place. Gray shading indicates the cochlear frequency region corresponding to the noise exposure band. **(D, E)** Ribbon volume range for IHCs **(D)** and OHCs **(E)** across low-(0.5–1.4 kHz), mid-(2–5.6 kHz), and high-frequency (8–32 kHz) cochlear regions. Black circles indicate controls, open red squares indicate 2 months post-exposure, and dark red squares indicate 10 months post-exposure. Ribbon volume ranges were estimated using a jackknife procedure. Error bars represent ±1 SD.

Mean IHC synapse counts did not differ significantly from controls at either post-exposure time point, indicating no persistent group-level loss of afferent synapses (Fig. 8B). OHC ribbon counts were reduced at 2 months post-exposure but did not differ from controls by 9–10 months, suggesting recovery of OHC presynaptic ribbon counts over time (Fig. 8C). In contrast, IHC and OHC ribbon volume ranges were larger at both 2 and 9–10 months post-exposure than in controls, indicating broader ribbon size distributions and a greater proportion of enlarged ribbons (Fig. 8D–E). Together, these findings indicate that although IHC synapse counts were preserved and OHC ribbon counts recovered after TTS, presynaptic ribbon-volume variability remained elevated.

Comprehensive histological analyses, including individual-ear variability, full cochlear ribbon volume distributions, and detailed analyses of hair cell survival, efferent innervation density, and ribbon morphology, are presented in Mondul et al. (2026). The following analyses therefore examined whether increased presynaptic ribbon-volume variability was associated with suprathreshold ABR measures.

#### 3.4.2 Inner hair cell ribbon synapse structure is not consistently associated with amplitude-based suprathreshold ABR measures

Across post-exposure time points and stimulus conditions, inner hair cell synaptic structure was not consistently associated with suprathreshold ABR amplitude measures. Analyses were performed at 80 dB SPL because this level yielded measurable Wave I responses in the greatest number of ears across frequencies and stimulus conditions. Synapse counts per inner hair cell were not associated with Wave I amplitudes for 80 dB SPL tonebursts of corresponding frequency at either post-exposure time point (2 months: β = 0.42, p = 0.841; 8–10 months: β = 1.41, p = 0.453), and no significant interaction with time point was observed (p = 0.445) (Fig. 9A). Presynaptic ribbon-volume variability, which increased following TTS, was not associated with Wave I toneburst amplitude at 2 months (β = 22.12, p = 0.557) but showed a negative association at 8–10 months post-exposure (β = −24.63, p = 0.025). However, the ribbon variability-by-time point interaction was not significant (Δβ = −40.37, p = 0.140), providing insufficient evidence that the strength of this relationship changed over time (Fig. 9B). Broadband click- and chirp-evoked Wave I amplitudes showed no consistent relationships with presynaptic ribbon-volume variability.

**Figure 9.**
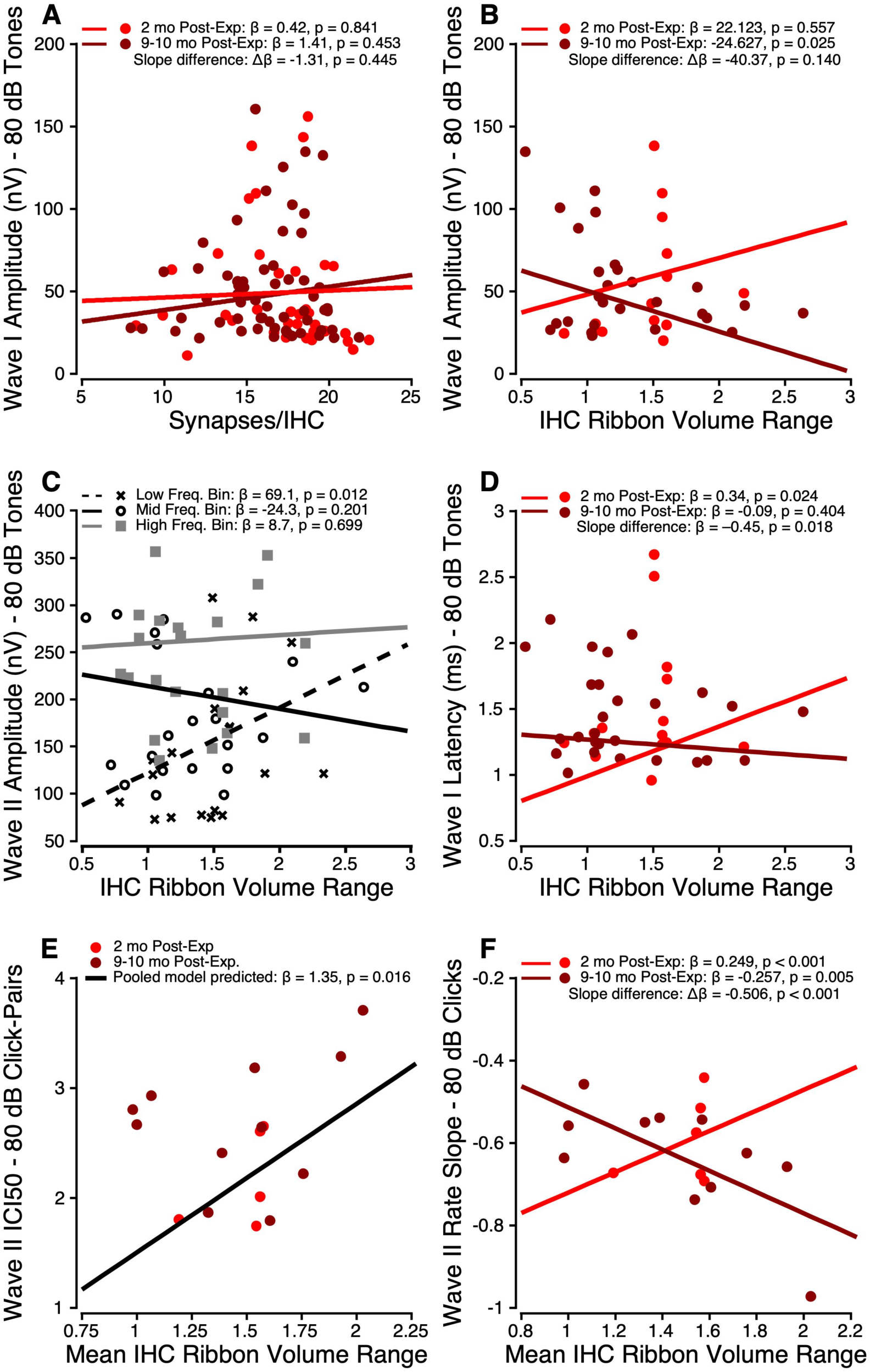
Relationships between inner hair cell synaptic structure and suprathreshold ABR measures. **(A)** Synapse count per inner hair cell versus Wave I amplitude (80 dB SPL tones) at 2 months (β = 0.42, *p* = 0.841) and 8–10 months (β = 1.41, *p* = 0.453); slopes did not differ (Δβ = −1.31, *p* = 0.445). **(B)** Ribbon volume variability versus Wave I amplitude at 2 months (β = 22.12, *p* = 0.557) and 8–10 months (β = −24.62, *p* = 0.025); slopes did not differ (Δβ = −40.37, *p* = 0.140). **(C)** Ribbon volume variability versus Wave II amplitude across frequency bins. Low-frequency: β = 69.1, *p* = 0.012; mid-frequency: β = −24.3, *p* = 0.201; high-frequency: β = 8.7, *p* = 0.699. The mid-frequency slope differed from the low-frequency slope (Δβ = −93.4, *p* = 0.003), whereas the high-frequency slope did not (Δβ = −60.5, *p* = 0.076). **(D)** Ribbon volume variability versus Wave I latency at 2 months (β = 0.34, *p* = 0.024) and 8–10 months (β = −0.09, *p* = 0.404); slopes differed (Δβ = −0.45, *p* = 0.018). **(E)** Mean ribbon volume variability versus Wave II ICI50 across post-exposure time points (β = 1.35, *p* = 0.016); no timepoint interaction (*p* = 0.224). **(F)** Mean ribbon volume variability versus Wave II rate-adaptation slope at 2 months (β = 0.249, *p* < 0.001) and 8–10 months (β = −0.257, *p* = 0.005); slopes differed (Δβ = −0.506, *p* < 0.001). Points represent individual ears; lines represent model-predicted fits.

Given the robustness of Wave II in macaque ABRs, we additionally evaluated structure– function relationships for Wave II amplitude. Presynaptic ribbon-volume variability was not associated with toneburst-evoked Wave II amplitude when frequencies were pooled, and this relationship did not differ across post-exposure time points (*p* = 0.983). However, frequency-bin analyses revealed region-specific relationships. Greater ribbon size variability was associated with larger Wave II amplitudes in the low-frequency bin (0.5–1.4 kHz; β = 69.1, *p* = 0.012). In contrast, the slope in the mid-frequency exposure region (2–5.6 kHz; β = −24.3, *p* = 0.201) was significantly more negative than the low-frequency slope (Δβ = −93.4, *p* = 0.003). No significant association was observed in the high-frequency bin (8–32 kHz) (Fig. 9C). Presynaptic ribbon-volume variability was also not consistently associated with broadband Wave II responses evoked by clicks or chirps at 80 dB SPL.

#### 3.4.3 Presynaptic ribbon-volume variability is associated with temporal ABR measures

In contrast to amplitude-based measures, ABR timing metrics revealed more consistent structure–function relationships. Synapse counts per inner hair cell were not consistently associated with suprathreshold temporal ABR measures, including Wave I or Wave II toneburst latency, paired-click ICI50, or rapid-rate click adaptation slope. Subsequent analyses therefore focused on presynaptic ribbon-volume variability, which remained elevated after TTS. At 2 months post-exposure, greater inner hair cell presynaptic ribbon-volume variability was associated with prolonged Wave I toneburst latency at 80 dB SPL (β = 0.34, p = 0.024), whereas the slope at 8–10 months was not significantly different from zero (β = −0.09, p = 0.404). The relationship differed significantly between post-exposure time points (Δβ = −0.45, p = 0.018), indicating that the association changed over time (Fig. 9D). The ribbon variability–latency relationship did not differ across frequency bins (*p* > 0.18), indicating no frequency-specific dependence, and was not observed for Wave II latency. Broadband latency measures evoked by clicks and chirps at 80 dB SPL likewise showed no consistent relationships with presynaptic ribbon-volume variability.

Additional suprathreshold ABR metrics of temporal processing, including paired-click and rapid-rate click paradigms at 80 dB SPL, further demonstrated time-dependent relationships with presynaptic ribbon-volume variability. Increased presynaptic ribbon-volume variability was associated with elevated Wave II ICI50 values (i.e., poorer temporal processing) across post-exposure time points (β = 1.35, p = 0.016), with no significant interaction with time point (p = 0.224) (Fig. 9E). In contrast, presynaptic ribbon-volume variability showed a time-dependent relationship with Wave II rate-adaptation slope. Greater ribbon size variability was associated with a more positive adaptation slope at 2 months post-exposure (β = 0.249, p < 0.001), but with a more negative adaptation slope at 8–10 months (β = −0.257, p = 0.005). The relationship differed significantly between post-exposure time points (Δβ = −0.506, p < 0.001) (Fig. 9F).

Together, these results indicate that timing-based ABR measures are more strongly associated with IHC synaptic remodeling following noise-induced TTS than amplitude-based ABR metrics.

## 4. DISCUSSION

This study demonstrates that moderate noise exposure sufficient to induce TTS produces long-lasting alterations in auditory brainstem processing that are not predicted by recovery of peripheral auditory measures such as DPOAEs and ABR thresholds. In a translational nonhuman primate model, peripheral auditory sensitivity recovered fully after noise exposure, whereas suprathreshold neural responses exhibited complex and dynamic changes in gain, synchrony, and temporal adaptability. Collectively, these findings reveal a dissociation between response magnitude and response fidelity: evoked responses were frequently preserved or enhanced, whereas neural synchrony, recovery, and adaptation remained impaired. This pattern extends prior behavioral, electrophysiological, and histological work in the same animals (Mackey et al., 2026; Mondul et al., 2026), providing converging evidence that acute noise exposures can produce sustained auditory dysfunction, even in the absence of cochlear hair cell or synapse loss.

### 4.1 Noise-induced TTS is associated with persistent suprathreshold temporal processing deficits despite recovered peripheral sensitivity and preserved or enhanced ABR amplitudes

Following noise-induced TTS, measures of peripheral auditory function normalized over time, including full recovery of DPOAEs and ABR thresholds. These physiological changes were accompanied by a lack of significant group-level synapse loss at both 2 and 9–10 months post-exposure (Mondul et al. 2026). In contrast, presynaptic ribbon volume distributions were broadened at both time points, indicating a persistent increase in heterogeneity of synaptic structure, consistent with reports of ribbon hypertrophy and expanded ribbon volume distributions following noise exposure (Hickman et al., 2020; Valero et al., 2017) and in age-related synaptopathy in other species (Liberman & Liberman, 2015). Such structural remodeling of ribbon synapses has been linked to altered vesicle pool organization and neurotransmitter release dynamics at IHC synapses (Matthews & Fuchs, 2010; Moser et al., 2006). Thus, despite recovery of peripheral auditory sensitivity and the absence of significant group-level synapse loss, noise-exposed macaques exhibited persistent changes in synaptic ribbon structure, accompanied by substantial alterations in suprathreshold auditory neural function.

Under conditions of low temporal demand, suprathreshold ABRs were preserved or enhanced following noise exposure, with click- and tone-evoked responses showing larger amplitudes across multiple waves, particularly at moderate-to-high stimulus levels and at later post-exposure time points. A similar suprathreshold enhancement has been reported in synaptopathic mice and has been interpreted as compensatory gain or altered synaptic efficacy (Suthakar & Liberman, 2021). Importantly, in the present study, these effects occurred despite preserved synapse counts, indicating that gain changes are not driven solely by deafferentation but instead may reflect adaptive plasticity within peripheral synapses and/or central auditory circuits that maintains response magnitude when temporal demands are minimal. Persistent ribbon hypertrophy may increase the readily releasable vesicle pool at surviving synapses, such that a given stimulus level elicits greater neurotransmitter release and larger-than-typical suprathreshold responses, even in the absence of synapse loss (Matthews & Fuchs, 2010; Wichmann & Moser, 2015).

Interestingly, evidence of suprathreshold enhancement was most apparent outside the primary exposure region. In addition to enhanced click- and tone-evoked ABRs, low-frequency DPOAE amplitudes and thresholds recovered beyond baseline at later post-exposure time points, and greater presynaptic ribbon-volume variability was associated with larger Wave II amplitudes in the low-frequency cochlear region. Although the mechanism underlying these low-frequency changes remains uncertain, their consistency across physiological measures suggests that long-term compensatory plasticity following moderate noise exposure may extend beyond the directly exposed cochlear region. Rather than reflecting improved auditory function, the low-frequency enhancement may represent homeostatic compensation that preserves or increases response magnitude despite persistent synaptic remodeling and impaired temporal fidelity (Auerbach et al., 2014; Chambers et al., 2016; Salvi et al., 2000).

Chirp-evoked ABRs exhibited a pattern distinct from that of clicks and tones, with reduced amplitudes and a uniform prolongation of latency, despite preserved level dependence. Because chirps are designed to maximize neural synchrony by compensating for cochlear traveling-wave delays (Elberling et al., 2007), these findings indicate impaired neural firing coordination under optimized timing conditions. This dissociation parallels observations of reduced binaural interaction components (BICs) of the ABR and impaired spatial hearing in the same animals (Mackey et al., 2026), suggesting that compensatory gain preserves response magnitude but does not restore precise temporal alignment across distributed neural populations. Similar reductions in synchrony-dependent responses have been reported in other species following noise exposure and with aging even when audiometric thresholds are preserved (Bharadwaj et al., 2015; Parthasarathy & Kujawa, 2018), highlighting temporal coordination as a vulnerable dimension of auditory processing.

Noise-exposed macaques exhibited impaired neural adaptation under increasing temporal demands, but these effects were only apparent from normalized ABR metrics. In both rapid-rate click and paired-click paradigms, raw responses were enhanced at slow presentation rates and long inter-click intervals, giving the appearance of preserved function. However, normalization revealed that responses declined more steeply with increasing repetition rate and recovered less effectively at shorter inter-click intervals. These seemingly contrasting findings, namely enhanced responses when temporal demands were low but poorer response preservation when temporal demands increased, have been reported in humans and animal models following noise exposure and with aging, where deficits emerge only when temporal precision is taxed (Bharadwaj et al., 2015; Fujihira et al., 2026; Parthasarathy et al., 2019; Shaheen et al., 2015).

Rather than reflecting improved neural function, enhanced suprathreshold responses therefore appear to coexist with persistent deficits in temporal fidelity, suggesting that compensatory gain and synaptic remodeling are parallel consequences of noise-induced TTS.

In summary, noise-induced TTS in macaques produced a persistent dissociation between response magnitude and response fidelity. Suprathreshold response amplitudes were preserved or enhanced under conditions of low temporal demand, whereas neural synchrony, recovery, and adaptation remained impaired through 9–10 months post-exposure.

### 4.2 Implications for ABR assays of cochlear synaptopathy

Noise-induced synaptopathy has been proposed as a primary mechanism underlying hidden auditory dysfunction. The present findings support the broader concept that noise exposure can produce hidden auditory dysfunction, showing substantial suprathreshold functional deficits in the absence of hair cell loss or permanent threshold shifts. Our macaques did not exhibit the classic loss of afferent ribbon synapses, but instead showed persistent ribbon hypertrophy (Mondul et al., 2026), suggesting altered synaptic function rather than simple loss of afferent input. Synaptic hypertrophy has been associated with altered glutamatergic release dynamics and enhanced onset responses (Matthews & Fuchs, 2010; Sheets et al., 2017; Wichmann & Moser, 2015), which could explain preserved or exaggerated suprathreshold responses under low temporal demand while simultaneously degrading sustained temporal integration, neural synchrony, and adaptability under more demanding conditions. Importantly, such synaptic or auditory nerve dysfunction may not be indexed by threshold-based measures, DPOAEs, or ribbon synapse counts alone and therefore provides a plausible substrate for the persistent suprathreshold temporal deficits observed here and in related behavioral and electrophysiological measures in these macaques (Mackey et al., 2026).

The structure–function analyses further support the idea that ribbon remodeling, rather than ribbon synapse loss, is associated with persistent alterations in suprathreshold auditory processing following TTS. Synapse counts were not consistently associated with suprathreshold ABR amplitudes or temporal measures, whereas presynaptic ribbon volume variability showed stronger and more consistent relationships with timing-based ABR metrics than with amplitude-based responses. Rather than implying that ribbon variability directly determines physiological response magnitude or timing, these findings suggest that increased ribbon size variability may serve as a marker of the degree of synaptic remodeling and associated central auditory changes following TTS. Ears exhibiting greater ribbon remodeling also exhibited altered suprathreshold auditory processing, although the direction and strength of these relationships depended on the physiological measure and post-exposure interval. Ribbon volume variability was associated with enhanced Wave II amplitudes only in the low-frequency cochlear region, whereas relationships with temporal measures, including Wave I latency, Wave II paired-click recovery, and Wave II rapid-rate click adaptation, were more robust across analyses. Together, these findings suggest that synaptic remodeling following TTS may coexist with compensatory enhancement of response magnitude under low temporal demand while preferentially disrupting the precision and adaptability of auditory signaling (Goutman & Glowatzki, 2007; Liberman et al., 2011; Matthews & Fuchs, 2010; Wichmann & Moser, 2015).

Clinically, recovery of behavioral or ABR thresholds and DPOAEs is often interpreted as resolution of auditory injury. However, the present findings demonstrate that substantial deficits in suprathreshold temporal processing persist beyond apparent peripheral recovery and are revealed only under conditions that tax neural adaptability. Similarly, recent behavioral work demonstrated that excitotoxic cochlear synaptopathy selectively impaired perception of brief acoustic onset cues while sparing detection of longer-duration signals, reinforcing the concept that hidden auditory dysfunction is preferentially revealed by tasks requiring precise temporal processing rather than simple detection (Loftus et al., 2026). Rate- and interval-based ABR paradigms, particularly when combined with normalization approaches, therefore provide sensitive assays of temporal fidelity that are not captured by conventional threshold-based diagnostics. Because similar ABR components and spatial hearing measures are conserved across macaques and humans (Laumen et al., 2016; Peacock et al., 2021), these approaches may offer translationally relevant biomarkers for detecting noise-induced auditory dysfunction that remains hidden in standard clinical testing.

## 5. CONCLUSION

A single moderate noise exposure sufficient to induce TTS produced persistent suprathreshold auditory dysfunction despite recovery of peripheral sensitivity and the absence of cochlear hair cell or synapse loss in a translational macaque model. Noise exposure produced a sustained dissociation between response magnitude and fidelity: suprathreshold responses were often preserved or enhanced under low temporal demand, whereas neural synchrony, recovery, and adaptation remained impaired. Presynaptic ribbon-volume variability may serve as a structural marker of the synaptic remodeling accompanying these functional changes, rather than a direct determinant of response amplitude. These findings support temporally demanding ABR paradigms, particularly rate- and interval-based measures, as translational biomarkers of auditory dysfunction that are hidden by threshold- and amplitude-based assessments.

## CRediT Author Contributions

RR and TAH designed experiments and secured funding. ANC, JAM, TAH, and RR conceptualized the analysis. ANC, JAM, SK, CAM, AB, and NT collected data. ANC, JAM, SK, and AB analyzed data. ANC wrote the paper. All authors revised the paper.

## Acknowledgements/Funding

The authors are especially grateful to Mary Feurtado for her expertise and assistance with anesthesia and to Drs. M. Charles Liberman and Leslie Liberman for their expertise and assistance in the processing and analysis of histological data. The authors also thank Katy Alek, Alex Tarabillo, Jessica Feller, Rachel Archer, David Pitchford, and Oscar Rausis for assistance with data collection and analysis. This work was supported by NIH–NIDCD R01 DC015988 (MPI: R. Ramachandran and B. Shinn-Cunningham).

